# Rapid High-Resolution Analysis of Polysaccharide-Lignin Interactions via Proton-Detected Solid-State NMR with Application to Eucalyptus and Spruce

**DOI:** 10.1101/2025.04.06.647447

**Authors:** Peng Xiao, Jayasubba Reddy Yarava, Debkumar Debnath, Priya Sahu, Yifan Xu, Li Xie, Daniel Holmes, Tuo Wang

## Abstract

The plant secondary cell wall, a complex matrix composed of cellulose, hemicellulose, and lignin, is crucial for the mechanical strength and water-proofing properties of plant tissues, and serves as a primary source of biomass for biorenewable energy and biomaterials. Structural analysis of these polymers and their interactions within the secondary cell wall has been heavily relying on ^13^C-based solid-state NMR techniques. In this study, we explore the application of ^1^H-detected solid-state NMR techniques for rapid, high-resolution structural characterization of polysaccharides and lignin, demonstrated on the stems of hardwood eucalyptus and softwood spruce. We employed several strategies, including the use of synthesized 2D spectra to resolve central ^1^H resonances and the combined application of 3D hCCH and hCHH experiments for complete resonance assignment and unambiguous identification of lignin-carbohydrate interactions. Our findings emphasize the central role of acetylated three-fold xylan conformers, rather than two-fold, in stabilizing the carbohydrate-lignin interface, with glucuronic acid sidechains in eucalyptus glucuronoxylan colocalizing with lignin, revised cellulose-lignin interactions involving uncoated microfibril surfaces, and pectin-lignin interactions indicative of early-stage lignification. We also carefully evaluated the interference from tannins in softwood species. These results present a novel approach for rapid structural analysis of lignocellulosic biomaterials without the need for solubilization or extraction.

## INTRODUCTION

The plant secondary cell wall is a chemically complex and structurally heterogeneous matrix that provides mechanical reinforcement, hydrophobicity, and resistance to biotic and abiotic stresses^1,2^. It is synthesized following the cessation of cell expansion, forming an intricate network primarily composed of three types of biopolymers. Cellulose microfibrils, composed of β-1,4-glucan chains hydrogen-bonded together, provide structural rigidity to the cell wall, while hemicelluloses, including xylan and mannan, exhibit species-specific variations in their backbone substitutions, which influence their physicochemical properties and interactions^3, 4^. Lignin, a polyphenolic polymer synthesized through the oxidative polymerization of monolignols, reinforces structural rigidity, reduces wall permeability, and improves microbial resistance^5-8^. The complex supramolecular organization and extensive interactions between these biopolymers dictate the functional properties of the secondary cell wall, significantly influencing plant biomechanics, water transport efficiency, and degradation resistance^9^. A detailed understanding of these interactions is critical for advancements in biofuel production, lignin valorization, and the development of sustainable biomaterials^10-12^.

Recently, multidimensional ^13^C solid-state NMR spectroscopy has emerged as a powerful tool for analyzing the molecular composition and physical interactions within secondary plant cell walls ^13-16^. This technique enables the study of natively hydrated plant tissues without the need for solubilization or extraction, preserving the native physical and chemical state of the biopolymers. As a result, molecular architectural models of secondary cell walls across a wide range of plant species have been proposed^17-20^. A key structural insight gained from these studies is the role of xylan substitution patterns and conformational structures in mediating the interactions between cellulose and lignin. Specifically, the flat-ribbon structure of xylan, stabilized by an evenly distributed pattern of substitutions, is essential for binding to cellulose microfibrils, while non-flat conformations promote interactions with lignin^21-24^. Furthermore, it was found that xylan serves as the primary interactor with lignin, with extensive electrostatic interactions between their polar functional groups, while cellulose interacts with lignin only as a secondary site^23, 25^. The specific roles of different monolignol units have also been more clearly defined, with the guaiacyl (G) unit being deposited early to interact with methylated pectin during the initial stages of lignification, while the methoxyl-rich syringyl (S) unit is deposited later, stabilizing the carbohydrate-lignin interface through extensive interactions with acetylated xylan^26^. These studies rely heavily on 2D/3D ^13^C correlation experiments on uniformly ^13^C-labeled plant tissues, providing high-resolution insights into the interaction interfaces of the three biopolymers within lignocellulosic materials.

Despite these significant advancements, several aspects of solid-state NMR techniques and their application to lignocellulosic biomass characterization remain open to further refinement^27^. One of the primary limitations is the inherently low sensitivity of NMR spectroscopy, a challenge that has been partially addressed through sensitivity-enhancing strategies such as magic-angle spinning dynamic nuclear polarization (MAS-DNP) and the development of solid-state cryoprobes^28-32^. Another major constraint is the requirement for isotopic enrichment, particularly ^13^C labeling, which, while costly, is being optimized through more economical labeling strategies and instrumentation^33, 34^. MAS-DNP has further mitigated this limitation by enabling the acquisition of 2D ^13^C-^13^C correlation spectra using the naturally present ^13^C (1%) in unlabeled samples, although this comes at the expense of spectral resolution due to the required cryogenic temperatures, particularly for soft matrix polymers^35-41^. Finally, insufficient spectral resolution hinders the differentiation of complex carbohydrate-lignin networks with diverse linkages, conformations, and spatial distributions, a challenge being addressed through the development of optimized experiments for carbohydrate polymers.

Another potential approach to overcoming the three bottlenecks is the use of ^1^H-detection solid-state NMR, which offers high natural isotopic abundance, high sensitivity, and the potential to enhance spectral resolution when combined with ^13^C and other nuclei^42-44^. This technique also allows for the use of smaller sample sizes, typically ranging from 1-10 mg, due to the use of smaller rotors (e.g., 0.7-1.3 mm) and the requirement for fast and ultrafast magic-angle spinning (MAS) at 60-110 kHz. Such methods are well established in protein studies, where a comprehensive toolbox for resonance assignment and structural analysis exists^45-48^. ^1^H-detection solid-state NMR has also been widely applied to pharmaceutical compounds for rapid quantification of active pharmaceutical ingredients (APIs), structural analysis in formulated pharmaceuticals, and monitoring the presence of residual water ^49-51^. Recently, efforts have adapted these methods to investigate the structure of cellular carbohydrates. Applications have been made to plant primary cell walls, enabling selective analysis of dynamic and semi-dynamic pectin and hemicellulose, cellulose, specifically analyzing hydroxymethyl conformations, as well as cell walls and capsules of both pathogenic and edible fungal species, and bacterial peptidoglycans^52-60^. This study explores the feasibility of using ^1^H-detection approaches to analyze the structure of carbohydrates and lignin, and to better understand their interactions within intact plant secondary cell walls.

## EXPERIMENTAL SECTION

Uniformly ^13^C-labeled mature stems of two plants, eucalyptus (*Eucalyptus grandis*) and spruce (*Picea abies*), were grown in ^13^C-enriched (97 atom%) closed-atmosphere chambers for 16 weeks at IsoLife (Wageningen, The Netherlands)^25^. The debarked plant stems were finely cut into small pieces using a razor, and packed into a 1.3 mm MAS rotor for solid-state NMR analysis. All solid-state NMR experiments were performed on a Bruker Avance-NEO 600 MHz (14.1 T) spectrometer equipped with a 1.3 mm triple-resonance HCN probe, operating at a MAS rate of 60 kHz. ^13^C chemical shifts were externally referenced using the tetramethylsilane (TMS) scale, with adamantane methylene resonance set at 38.48 ppm, while ^1^H chemical shifts were referenced to sodium trimethylsilylpropanesulfonate (DSS) at 0 ppm. The cooling gas temperature was maintained at 250 K, with an estimated sample temperature of approximately 296 K, accounting for heating effects from fast MAS.

Short-range (predominantly one-bond) ^1^H-^13^C correlations were acquired using a 2D hCH experiment with a short second CP contact time, which was set to 100 μs for eucalyptus and 50 μs for spruce. Through-bond ^13^C-^13^C connectivity was established using a 3D hCCH-TOCSY (total correlation spectroscopy) experiment^46^, employing a 15-ms WALTZ-16 (wideband alternating-phase low-power technique for zero-residual splitting) mixing^61^ with a radiofrequency (rf) field strength of 21.4 kHz. Through-space ^1^H-^1^H interactions were probed in eucalyptus using 2D hChH and 3D hCHH experiments, incorporating RFDR-XY16 (radio frequency-driven recoupling) mixing sequences^62, 63^ with recoupling times ranging from 0.133 ms to 0.8 ms. For spruce, long-range interactions were analyzed using a 2D hCH experiment with a 1.2-ms second CP contact time. Water suppression for all experiments was achieved using the MISSISSIPPI (Multiple Intense Solvent Suppression Intended for Sensitive Spectroscopic Investigation of Protonated Proteins) sequence^64^, with a 100 ms duration and an rf field strength of 15.4 kHz.

For the eucalyptus sample, heteronuclear dipolar decoupling was applied using slpTPPM (swept low-power two-pulse phase modulation) decoupling^65^ at 13.54 kHz rf power on the ^1^H channel during t_1_ evolution in hCH and 2D/3D hCHH RFDR sequences. WALTZ-16 decoupling at 10 kHz was applied on the ^13^C channel during direct detection of ^1^H chemical shift evolution in the 2D hCH, 2D hChH, and 3D hCHH experiments. No decoupling was applied during the indirection dimension of ^1^H chemical shift evolution, instead a π pulse was implemented for refocusing. For 3D hCCH TOCSY, slpTPPM decoupling was applied during t_1_ and t_2_ evolution on the ^1^H channel, while WALTZ-16 decoupling at 10 kHz rf power was applied on the ^13^C channel during direct ^1^H detection. The 90° pulse lengths were 2.5 μs for ^1^H (100 kHz rf power) and 5 μs for ^13^C (50 kHz rf power). ^1^H-^13^C cross-polarization (CP) was performed under the double-quantum (n = +1) Hartmann-Hahn condition, with rf powers of 49.675 kHz for ^1^H and 10 kHz for ^13^C, at an MAS rate of 60 kHz. For the spruce sample, slpTPPM decoupling was applied during t_1_ evolution with rf field strength of 13.54 kHz, while WALTZ-16 decoupling at 14.9 kHz rf power was applied on the ^13^C channel during ^1^H direct detection. 2D and 3D datasets were acquired using the States-TPPI method^66^. Detailed experimental parameters are provided in **Tables S1**, while the assigned ^13^C and ^1^H chemical shifts of carbohydrate and lignin polymers are listed in **Tables S2 and S3**.

Additional 2D CP refocused J-INADEQUATE spectrum of spruce was recorded on a 600 MHz (14.1 T) Bruker spectrometer equipped with a 3.2mm MAS probe operating at 14 kHz MAS^67, 68^. The J-evolution period was 1.42 ms for each of the four τ periods. The sample temperature was at approximately 294 K. The rf strengths were 83 kHz for ^1^H decoupling, and 83.3 kHz and 62.5 kHz for the 90° hard pulses of ^1^H and ^13^C, respectively.

## RESULTS AND DISCUSSION

### Synthesized hCH spectra reveal central ^1^H resonances of polysaccharides and lignin

A proton-detected 2D heteronuclear correlation hCH spectrum was acquired on the stem tissue of *Eucalyptus grandis*, revealing a collection of well-resolved signals originating from both cell wall polysaccharides and lignin. These ^13^C-^1^H cross peaks predominantly correspond to one-bond correlations, as indicated by the absence of signals from non-protonated syringyl (S) units at the S3/5 positions (153 ppm), which are two bonds away from the protonated S2/6 sites (**Figure 1A**). Among the polysaccharide components, cellulose signals were most distinctly resolved at the C4 positions of both interior and surface chains, exhibiting ^13^C chemical shifts of 89 ppm and 84 ppm, respectively (**Figure 1A**). Despite their shared chemical identity as β-1,4-glucan chains (**Figure 1B**), interior and surface cellulose differ in their hydrogen-bonding arrangements and conformational states. Interior chains exhibit double-sided hydrogen bonding and adopt a *trans-gauche* (*tg*) conformation of the exocyclic hydroxymethyl group, whereas surface chains engage in single-sided hydrogen bonding and primarily adopt a *gauche-trans* (*gt*) conformation, with a possible minor population of *gauche-gauche* (*gg*) conformers, resulting in distinct ^13^C4 chemical shifts^69, 70^. However, their corresponding ^1^H4 signals both centered near 3.5 ppm (**Figure 1A**). Due to the high proton density of the biomolecular matrix and broad conformational heterogeneity, these ^1^H resonances are extensively broadened, spanning from 2.1 to 4.8 ppm (**Figure 1A**).

**Figure 1.**
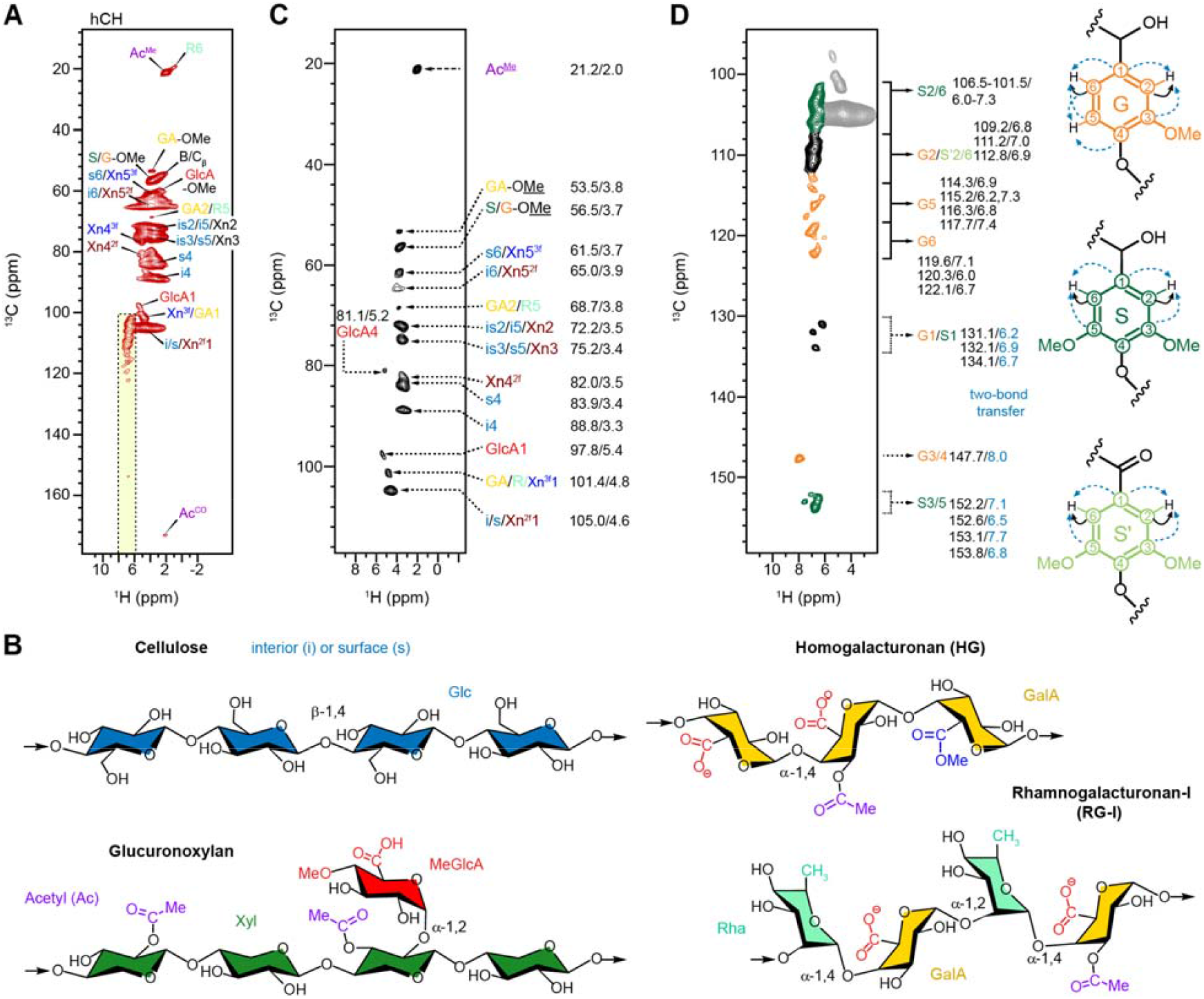
2D hCH spectra resolving various carbon and proton sites in carbohydrate and lignin. (**A**) hCH spectrum with a short 0.1 ms ^13^C-^1^H CP contact time showing carbohydrate carbons and their correlation with directly bonded protons. NMR abbreviations are used: surface cellulose: s; interior cellulose: i; two-fold xylan: Xn^2f^; three-fold xylan: Xn^3f^; galacturonic acid (GalA): GA; rhamnose (Rha): R; glucuronic acid: GlcA; Acetyl methyl and carbonyl: Ac^Me^ and Ac^CO^; syringyl: S; guaiacyl: G; methoxyl: OMe. Yellow strip highlights lignin aromatic ^1^H (6-8 ppm). (**B**) Representative structure of cellulose, hemicellulose and key pectin polymers in eucalyptus. (**C**) Synthesized 2D hCH spectrum using only the top contour levels of each peak regions for enhanced resolution on one-bond ^13^C-^1^H correlation. Each carbon site and the corresponding ^13^C/^1^H chemical shifts (ppm) are listed. (**D**) Synthesized 2D hCH spectrum showing the top contour levels of each lignin peak, with their ^13^C/^1^H chemical shifts (ppm) listed. Non-protonated sites (G1/S1, G3/4, S3/5) are displayed at adjusted contour levels to enhance visibility of weak 2-bond correlations. S units are shown in green, G units in orange, mixed S/G regions in black, and the carbohydrate region in grey. The structures of lignin units and observed ^13^C-^1^H polarization transfers are shown, with one-bond transfers indicated by black solid lines and two-bond transfers by light blue dashed lines.

To address ^1^H spectral complexity, an artificial spectrum was generated by extracting only the top contours of each peak, thereby enabling the identification of central ^1^H chemical shifts that represent the most probable structural populations (**Figure 1C**). Using this approach, the central ^1^H shifts for the C4 positions of interior and surface cellulose were determined to be 3.3 ppm and 3.4 ppm, respectively. A limitation of this contour-based method is the loss of fine structural resolution in the ^1^H dimension, particularly for highly polymorphic regions of cellulose. Specifically, 3-4 conformational variants of interior glucan chains and 2-3 distinct surface chain types—previously resolved in native plant cell wall cellulose due to hierarchical organization within and between microfibrils—are no longer discernible^71^. This highlights the need for further methodological improvements to resolve these polymorphic features in future studies, likely through partial deuteration or the use of ultrafast MAS techniques^59^.

The hemicellulose of the secondary cell wall in eucalyptus consists almost exclusively of glucuronoxylan, which is composed of a β-1,4-linked xylosyl backbone with glucuronic acid sidechains, which are often O-4 methylated, and acetyl substitutions (**Figure 1B**). Distinct signals corresponding to both two-fold and three-fold screw conformations of xylan were resolved, as evidenced by the Xn4^3f^, Xn4^2f^, and Xn1^3f^ resonances (**Figure 1A**). Glucuronoxylans in eucalyptus are heavily acetylated, with 50-70% of xylosyl residues bearing acetyl groups at the O2 and/or O3 positions^72-74^. The methyl group of the acetyl substituent (Ac^Me^) exhibited well-resolved signals at (21.2, 2.0 ppm) in the hCH spectrum (**Figure 1A, C**). The corresponding ^1^H resonance also showed a weak, two-bond correlation with the carbonyl carbon of the acetyl group (Ac^CO^) at 174 ppm, confirming its identity (**Figure 1A**).

The acetylation pattern of xylan can significantly alter the ^1^H chemical shifts of H2, H3, and H4, drifting between the 3.5-5.0 ppm range^75-78^. This contributes to the broad ^1^H signals of the Xn2 and Xn3 peaks (72.5 and 74.5 ppm), which further overlap with cellulose resonances, obscuring their ^1^H chemical shifts in standard 2D experiments and necessitating 3D techniques for resolution, as will be shown later in this study. Acetylation can also influence ^13^C chemical shifts, though to a lesser extent. For instance, the “mixed xylan” resonance previously observed in eucalyptus and poplar, which displays a C4 chemical shift near 80.5 ppm—intermediate between those of the two- and three-fold screw conformers—may also reflect heterogeneity in acetylation positions (and possibly MeGlcA substitutions as well), given its absence in spruce, a species in which xylan is not acetylated^25^.

Clear and unambiguous signals were detected for glucuronic acid (GlcA) residues. In particular, GlcA1 and GlcA4 resonances were observed at (97.8, 5.4 ppm) and (81.1, 5.2 ppm), respectively (**Figure 1A, C**). These arise from α-1,2-linked GlcA substituents at the O2 position of xylosyl residues, with a GlcA-to-Xyl molar ratio of approximately one-to-ten^79^. A distinct GlcA-OMe signal was observed at (59.6 ppm, 3.6 ppm) in **Figure 1A**, corresponding to the methoxy group of 4-O-methyl-glucuronic acid (MeGlcA), as shown in its molecular structure in **Figure 1B**^79, 80^. The identification of this signal—missed in previous ^13^C-based studies due to insufficient resolution—represents an opportunity in characterizing xylan branching patterns *in muro*.

The stem tissues analyzed contain a minor but detectable fraction of primary cell walls, formed prior to secondary wall deposition^81, 82^. Its presence is evident from the detection of galacturonic acid (GalA; GA) and rhamnose (Rha; R) signals—key components of pectic polysaccharides such as homogalacturonan and rhamnogalacturonan-I (**Figure 1B**)^83, 84^. Distinct peaks corresponding to GalA and Rha, including GA2/R5 at 68.7, 3.8 ppm and R6 at 19.4, 0.9 ppm, were observed (**Figure 1A, C**).

The lignin in Eucalyptus is predominantly composed of syringyl (S) units, with a syringyl-to-guaiacyl (S/G) ratio of approximately three-to-one^81^. Signals corresponding to S and its oxidized variant S′, a subpopulation featuring Cα oxidation (Cα=O), are best resolved through their distinct ^13^C resonances: the non-protonated S3/5 carbons at 152-154 ppm and the protonated S2/6 carbons at 101.5-107 ppm (**Figure 1D**). The ^1^H chemical shifts of S2/6 protons span the range of 6.0-7.3 ppm. Due to the non-protonated nature of S3/5, their cross-peaks in ^1^H-detected experiments rely on polarization transfer from remote protons, such as those on S2/6. Similar two-bond magnetization transfers were also observed for other non-protonated sites, including G1/S1 and G3/4, as demonstrated in the synthesized spectrum (**Figure 1D**). The synthesized spectrum serves as a useful tool not only for identifying the central ^1^H resonances of the most populated conformers within each carbohydrate but also for enabling resonance assignment of lignin in the solid state. The 2D hCH experiment, which typically takes 1-4 hours (**Table S1**), compared to 1-2 days for each ^13^C-based 2D experiment, enables rapid assessment of the structure of polymers.

### Tracking through-bond connectivity and full resonance assignment through 3D hCCH

The 3D hCCH TOCSY experiment, originally developed for protein sidechain assignments, was adapted to track carbon and proton connectivities in carbohydrates using J-coupling-based WALTZ-16 mixing^46, 55, 85^. This enabled unambiguous tracking of ^13^C chemical shifts for interior and surface cellulose in both ^13^C-^13^C (δ_1_-δ_2_) and ^13^C-^1^H (δ_1_-δ_3_) planes of the 3D hCCH spectrum (**Figure 2A**), a level of detail not achievable with 2D hCH experiments. Similarly, carbon sites in both two-fold and three-fold xylan conformations, as well as GlcA branches, were successfully tracked (**Figure 2B**). Notably, two types of GlcA residues were distinguished based on peak multiplicity at the C2/5 position, as well as at C4 and C1 positions. Two distinct forms of GalA units were also resolved based on their unique C1/H1 chemical shifts at (100.8, 5.1 ppm) and (101.4, 4.8 ppm), as well as differences at C4/H4 at (79.8, 4.7 ppm) and (79.0, 4.3 ppm), with another minor form appearing as a shoulder peak (**Figure 2C**). This may originate from the structural complexity of pectin, where GalA units can form the homopolymer HG, but with diverse patterns of methyl esterification and acetylation, alongside the presence in RG-I, where GalA coexists with Rha along the backbone^84^. The resolvable ^13^C chemical shifts from the analysis of 3D hCCH spectrum and the ^1^H chemical shifts from the 2D hCH spectra are summarized in **Figure 2D** and **Tables S2, S3**, representing a comprehensive resonance mapping of rigid cell wall biopolymers in fully protonated plant tissues using ^1^H-based solid-state NMR approaches.

**Figure 2.**
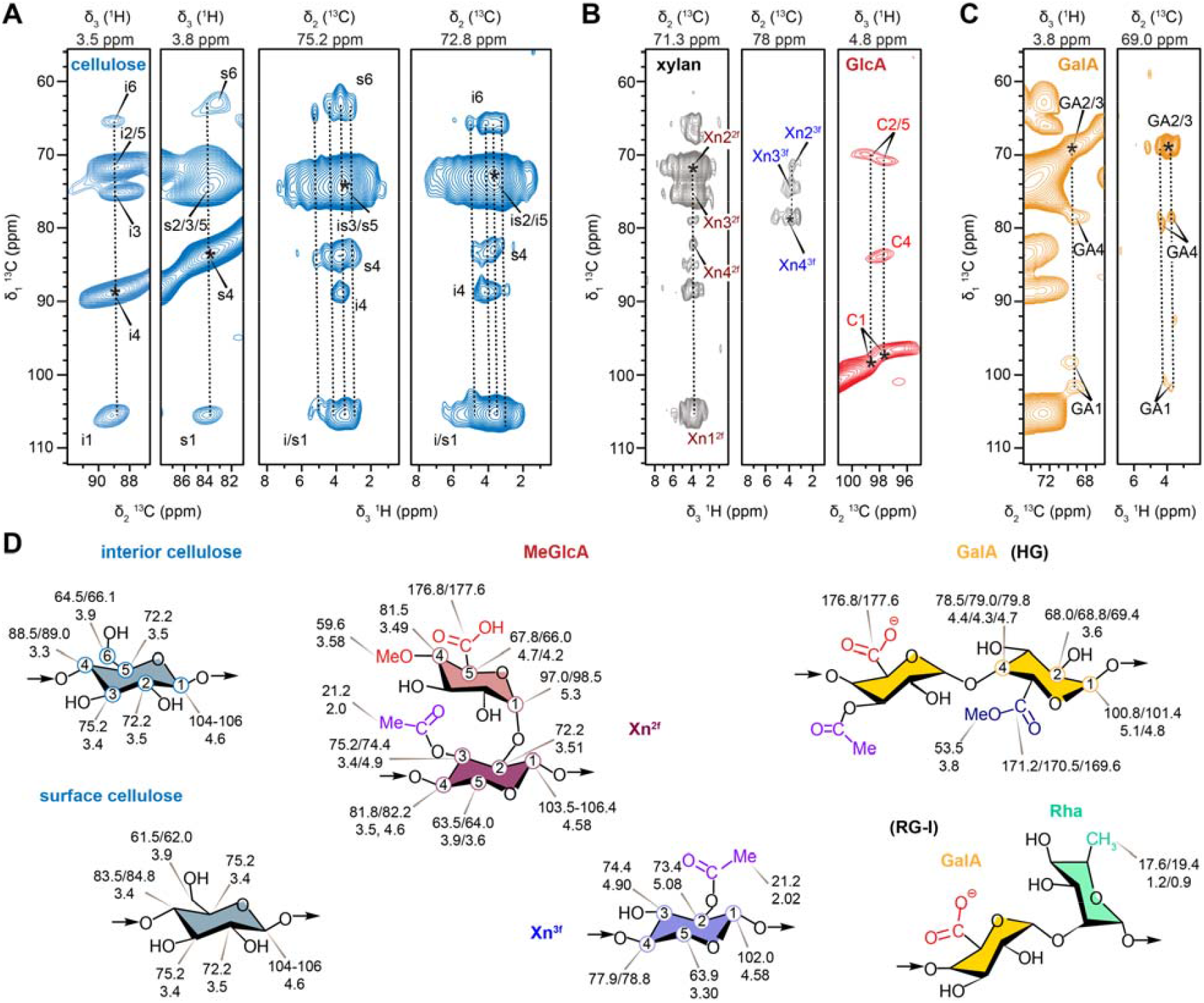
Through-bond connectivity within rigid polysaccharides of eucalyptus cell walls. 2D ^13^C-^13^C and ^13^C-^1^H strips of 3D hCCH spectra are shown for (**A**) cellulose, (**B**) hemicellulose, and (**C**) pectin. Through-bond connectivity was established using TOCSY experiment with a WALTZ-16 ^13^C-^13^C (δ_1_-δ_2_) mixing time of 15 ms, followed by a short 100 μs ^13^C-^1^H (δ_2_-δ_3_) CP. 2D planes were extracted at different carbon (δ_2_) and proton (δ_3_) dimensions (shown on the top of each panel) for resonance assignment. The diagonal peak is marked with an asterisk. The sequential carbon connectivity patterns are highlighted with vertical dashed lines. (**D**) Saccharide units in eucalyptus polysaccharides with resolved ^13^C (top) and ^1^H (bottom) chemical shifts (ppm) labeled for each carbon site.

### Mapping lignin-carbohydrate interactions via combined analysis of 3D hCCH and hCHH

To explore physical packing interactions between lignin and carbohydrates, we first tested the 2D hChH experiment with varying RFDR mixing periods to enhance polarization transfer between different polymers. Comparison with a standard hCH spectrum revealed new signals in the hChH spectra, including long-range correlations within the lignin network, such as S3/5-S/G^H^, S3/5-OMe^H^, and OMe-S3/5^H^ (**Figure 3A**). The dashed box in **Figure 3A** further highlights new interactions between carbohydrate carbons and lignin protons in the secondary plant cell walls. Additionally, a cross-peak between rhamnose C6 and acetyl methyl protons (R6-Ac^Me^) was observed, arising from the close spatial proximity of rhamnose and acetylated GalA units in the pectin matrix of primary plant cell walls (**Figure 2D**). Extending the RFDR period from 133 μs to 800 μs revealed stronger and more extensive intermolecular interactions (**Figure 3B**). Both the methyl and carbonyl carbons of acetyl groups (Ac^Me^ and Ac^CO^), primarily present in xylan and less so in pectin, exhibited strong cross-peaks with the S2/6 and G2/5/6 protons of lignin. Similarly, the S3/5 carbons of lignin showed cross-peaks with the acetyl methyl protons (Ac^Me^). These cross-peaks further underscore the essential role of acetyl groups and xylan in stabilizing the lignin-carbohydrate interface (**Fig. 3c**).

**Figure 3.**
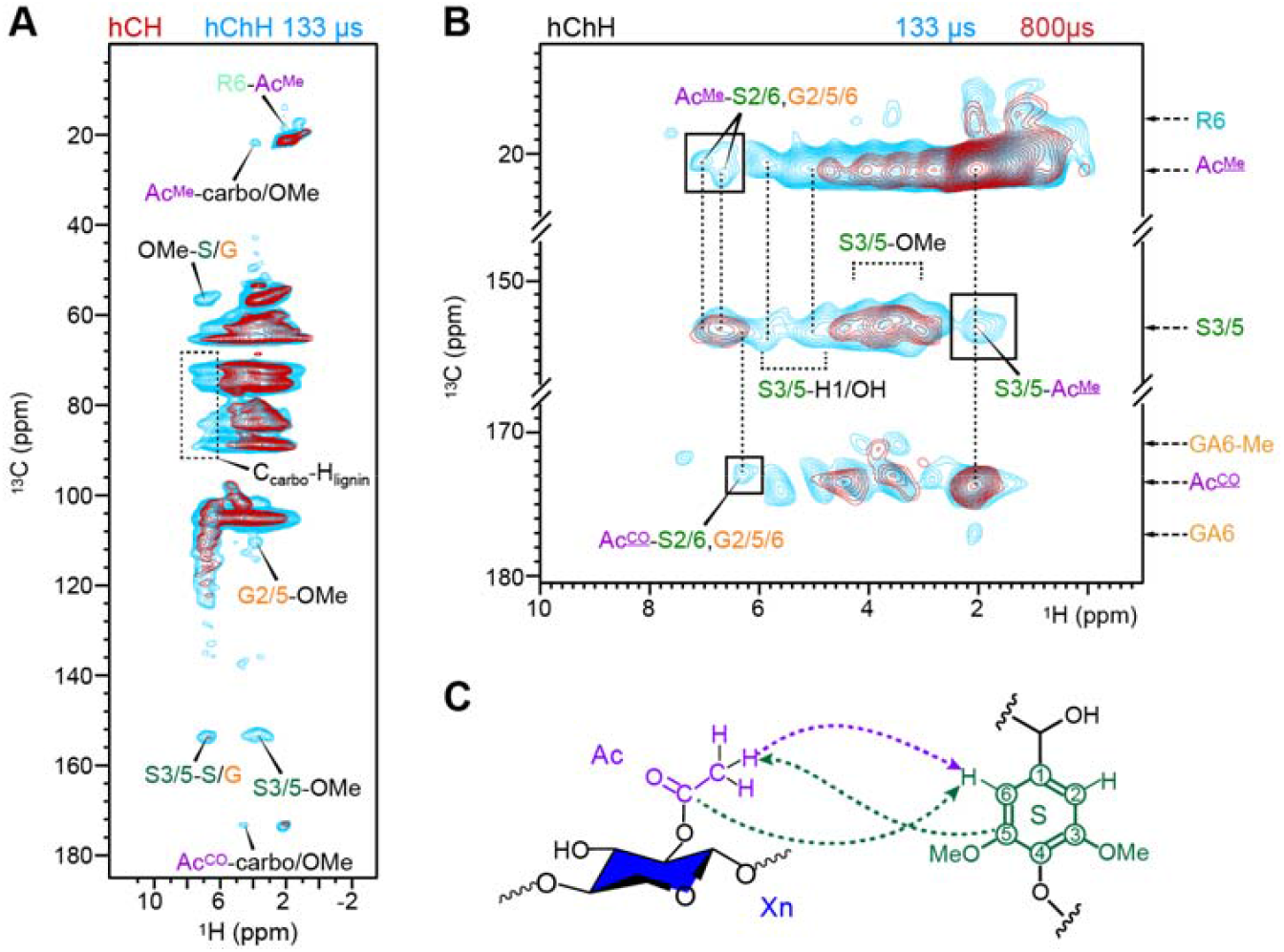
2D hChH probing lignin-carbohydrate intermolecular interactions. (**A**) Overlay of 2D hCH (dark red) and hChH spectra (cyan). The hChH spectrum was measured with 133 μs RFDR ^1^H-^1^H mixing, thus showing additional long-range interactions. (**B**) Selected regions of hChH spectra with varied RFDR mixing times. An 800 μs mixing (orange) enables detection of additional intermolecular interactions. Boxed regions highlight unambiguous intermolecular cross-peaks between the aromatic carbons and Ac protons and between the Ac carbons and aromatic protons. Certain interactions between S3/5 carbon sites and carbohydrate H1/OH protons (4.5-6.0 ppm, out of the range of OMe protons) can also be identified. Vertical dashed lines guide the comparison of key ^1^H sites. (**C**) Structural summary of observed interactions between xylan acetyl and lignin S-unit.

A more detailed, site-specific analysis of lignin-carbohydrate interactions is achieved by combining the hCCH TOCSY and hCHH RFDR spectra (**Figure 4**). The hCCH TOCSY spectrum resolves key ^13^C and ^1^H signals for a specific carbon site, while the hCHH RFDR spectrum extends to the ^1^H signals of all molecules spatially proximal to that original carbon site, thus providing a comprehensive view of intermolecular interactions. For example, three subforms of two-fold xylans (Xn4^2f^), which share a similar C4 signal at 82.2 ppm, were resolved based on their H4 signals in the 3.5-4.6 ppm range in the hCCH TOCSY spectrum (**Figure 4A**). All these subforms exhibited cross-peaks with lignin protons at 6.7 ppm in the hCHH RFDR spectrum (**Figure 4A**). At the same lignin 6.7 ppm ^1^H position, a weak cross-peak was also observed for Xn4^2f^/GlcA4 with a C4 signal at 81.4 ppm. The three-fold xylan, resolved at the 77.6 ppm plane, showed much stronger interactions with lignin aromatic protons at 6.6, 7.0, and 7.2 ppm (**Figure 4A**). In addition, cross-peaks observed at the Ac^Me^ site of xylan also matched these chemical shifts of lignin protons. These experimental evidences support the structural concept that three-fold xylan plays a more significant role than two-fold conformers in interacting with lignin.

**Figure 4.**
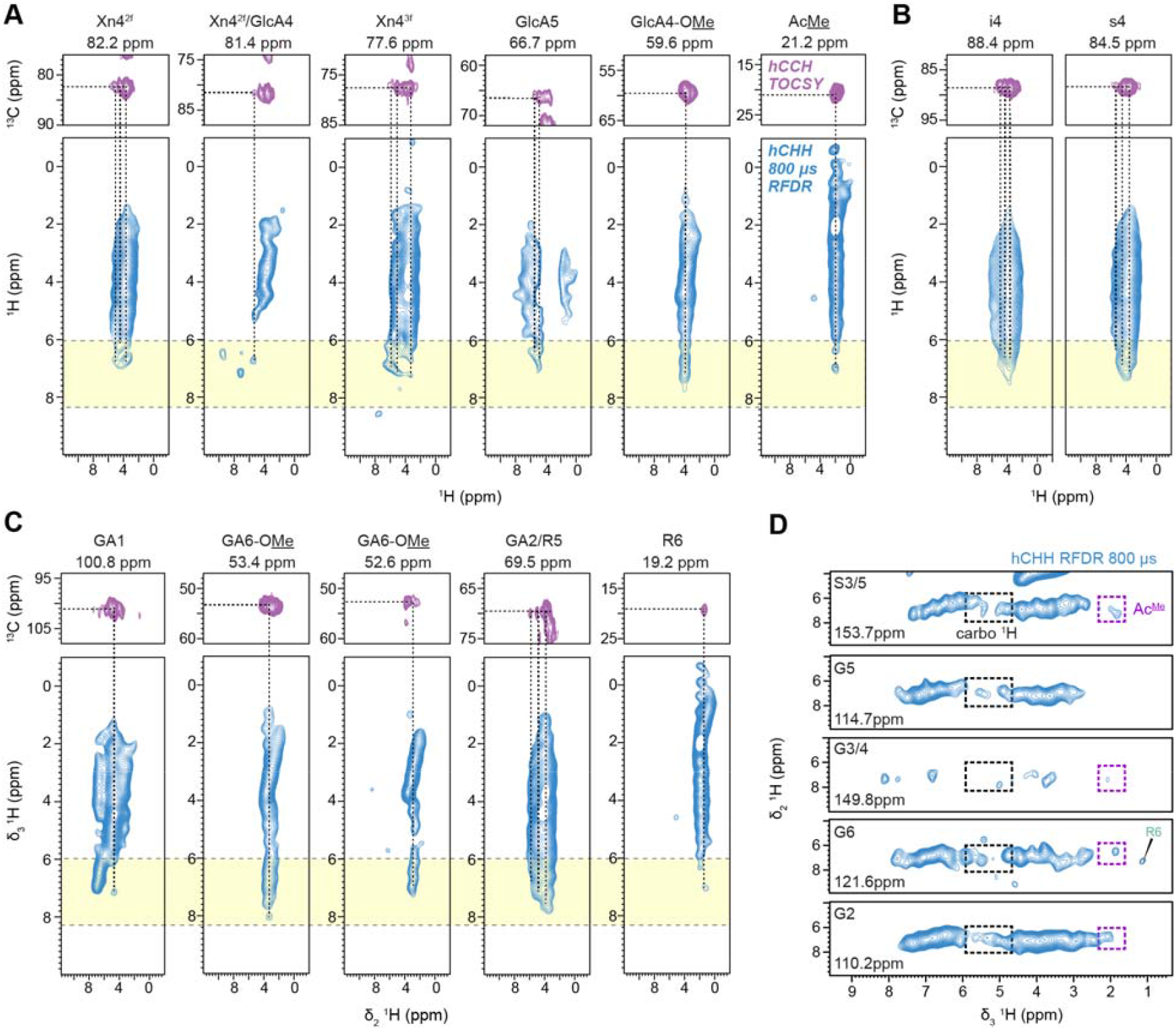
3D hCCH and hCHH spectra show intermolecular interactions with enhanced resolution. 2D strips extracted from 3D hCCH TOCSY (top, purple) and 3D hCHH RFDR (bottom, light blue) spectra are shown for (**A**) hemicellulose, (**B**) cellulose, and (**C**) pectin. The 2D planes from both 3D spectra share the same initial ^13^C site in the δ□ dimension, with the site and chemical shift labeled above each strip. The hCCH experiment, using 15 ms ^13^C-^13^C TOCSY mixing and 100 μs ^13^C-^1^H CP, shows intramolecular ^13^C-^13^C (δ_1_/δ_2_) and ^13^C-^1^H (δ_2_/δ_3_) correlations. Horizontal dashed lines mark ^13^C sites where ^1^H signals originate; vertical dashed lines indicate nearby ^1^H sites spatially proximal to the ^13^C sites. The hCHH experiment was carried out with 500 μs ^13^C-^1^H CP and 800 μs ^1^H-^1^H RFDR mixing, showing intramolecular ^13^C-^1^H (δ_1_/δ_2_) and intermolecular ^1^H-^1^H (δ_2_/δ_3_) interactions. Yellow boxes highlight the aromatic proton region in δ_3_ dimension and the cross-peaks in this box jointed by the dashed lines indicate unambiguous carbohydrate-lignin interactions. (**D**) 2D ^1^H-^1^H planes extracted from 3D hCHH spectrum showing cross peaks between lignin ^13^C and carbohydrate ^1^H. Each lignin ^13^C site (δ_1_) was indicated in the panels. The black and purple boxes highlight cross peaks of lignin protons (δ_2_) with unambiguous carbohydrate protons and Acetyle protons (δ_3_). A weak lignin-pectin cross-peak between G6 and R6 is also identified.

The strong cross-peaks with lignin protons observed at the GlcA4-OMe site (the 59.6 ppm plane) and GlcA5 (the 66.7 ppm plane) are unexpected (**Figure 4A**). While the methylated GlcA sidechains of glucuronoxylan have been proposed to form covalent linkages with lignin, mostly from model studies, no experimental evidence *in muro* has yet supported this hypothesis^4, 86^. These cross-peaks, however, indicate a close spatial proximity, or colocalization, of xylan’s GlcA sidechains with lignin.

The cellulose s4 and i4 sites also exhibited substantial cross-peaks with lignin protons—stronger than those observed for two-fold xylan’s interactions with lignin (**Figure 4B**). While it has been proposed that two-fold xylan coats the surface of cellulose, creating junctions that may serve as secondary interaction sites for lignin (following lignin’s primary interactions with three-fold xylan), the current ^1^H data suggest otherwise. Instead, these results imply that lignin may interact more extensively with regions of cellulose microfibrils that are not coated by two-fold xylan, possibly due to the lower abundance of two-fold xylan relative to cellulose.

A recent study has shown evidence of pectin’s interactions with lignin, particularly with G units, during the early stages of lignification^26^. In our hCHH RFDR spectra, pectin exhibited cross-peaks with lignin, with relatively weak, single cross-peaks observed at the GA1 and R6 sites (**Figure 4C**). In contrast, more extensive interactions were detected at the methyl ester sites (GA6-OMe) that appeared at two distinct ^13^C chemical shifts (53.4 and 52.6 ppm). This new observation indicate that methyl-esterified pectin is closely packed with lignin. Meanwhile, the GA2/R5 site at 69.5 ppm showed the strongest apparent interaction with lignin; however, this signal may result from overlap with GlcA resonances of xylan sidechain and is therefore not considered in this analysis.

While the previously discussed data rely on correlations extending from well-resolved carbohydrate C/H sites to lignin protons, an alternative approach extends from resolved lignin carbon sites to carbohydrate ^1^H signals (**Figure 4D**). The S3/5 sites displayed strong correlations with carbohydrate protons and acetyl methyl protons. Only regions free from overlap with lignin aromatic and methoxy proton signals are highlighted. Similar interactions were observed for carbon sites in G units, but these were generally weaker and, in some cases, absent—for example, the lack of an AcMe signal in the G5 plane. This pattern reinforces the stronger role of S-lignin in interacting with xylan.

In parallel, we also explored the analysis of δ_1-_δ_2_ ^13^C-^1^H planes extracted from the 3D hCHH spectrum (**Figure S1**). However, this approach necessitates slicing along the δ_3_ ^1^H dimension, which exhibits low resolution. As a result, the extracted planes frequently contain overlapping signals from both carbohydrate and lignin components, leading to ambiguity. Only a limited number of planes—specifically those corresponding to lignin ^1^H chemical shifts at 7.1 ppm and 6.6 ppm—displayed relatively reduced spectral congestion. This allowed identification of distinct cross-peaks for two-fold xylan and cellulose carbons with lignin protons, while signals from three-fold xylan remained fully overlapped and unresolvable. These results suggest that alternative experimental strategies, such as hCChH or methods incorporating spectral editing techniques to selectively suppress specific signal contributions, should be further investigated to enhance component-specific resolution in future studies.

### Resolving tannin and lignin signals in spruce

In the course of extending the ^1^H-detected solid-state NMR analysis from hardwood eucalyptus to softwood spruce, we encountered an unanticipated complication: significant spectral overlap between tannin-derived signals and those of G lignin units, which are the dominant monolignols in spruce lignin (**Figure 5A**). Nevertheless, three distinct resonances attributable to tannins were identifiable, particularly at the C4/H4 positions (T4), where at four structurally distinct environments could be resolved at (35.8, 4.95 ppm), (37.3, 4.4 ppm), (39.9, 2.0 ppm), and (41.4, 2.7). Additional tannin-derived signals, such as T6/8/10, were also distinguishable from those of carbohydrates and lignin^87^. Tannin signal assignment was aided by a 2D ^13^C refocused J-INADEQUATE experiment, which traces carbon-carbon connectivity and utilizes a double-quantum (DQ) dimension to enhance spectral dispersion and resolution, allowing us to distinguish ^13^C signals from G-lignin and tannin (**Figure 5B**)^67, 68^. Additionally, improved debarking procedures can be employed, if necessary, to remove the majority of tannins and minimize their interference with lignin analysis.

**Figure 5.**
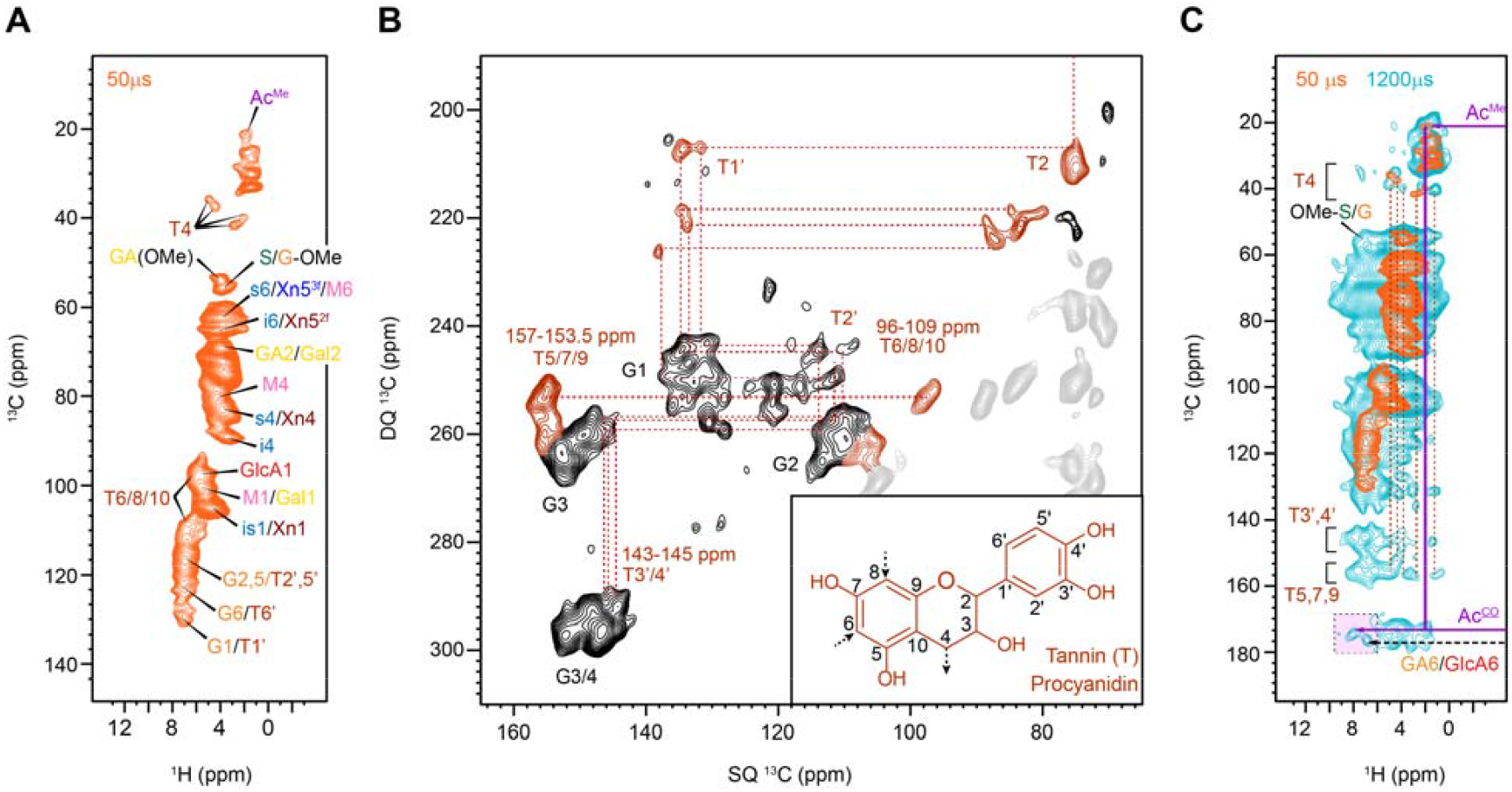
Analysis of carbohydrate and lignin components in spruce. (**A**) 2D hCH spectrum of spruce with a short ^13^C-^1^H CP contact time of 50 μs showing only one-bond correlations. (**B**) 2D CP ^13^C refocused J-INADEQUATE spectrum of spruce showing signals of G-lignin (back) and connectivity of tannin carbons (brown). Signature tannin peaks can be identified for T5/7/9 at 153-157 ppm, T6/8/10 at 96-109 ppm, and T3’/4’ at 143-145 ppm. (**C**) Overlay of two 2D hCH spectra with short (orange) and long (blue) CP contact times. Many tannin cross-peaks are observed and are indicated with brown vertical dashed lines. Carbohydrate-lignin cross-peaks are highlight in purple box. Connectivity of acetyl signals are highlighted using purple lines.

Simultaneously, we observed rigid-domain signals corresponding to two- and three-fold xylan conformations, internal and surface cellulose chains, and mannose (M) residues from galactoglucomannan—a hemicellulose prominently enriched in softwoods (**Figure 5A**). Acetyl group resonances (Ac^Me^) were detected as well; however, unlike in eucalyptus where acetylation predominantly occurs on xylan, these groups are now associated with galactoglucomannan in spruce secondary cell walls. In addition, spruce has arabinoglucuronoxylan with both arabinose and GlcA substitutions^4, 88^, but arabinoses are typically mobile and thus not observed in the hCH spectrum with a very short CP contact time^25^.

Extending the CP contact time in hCH spectrum from 50 μs to 1200 μs allowed tannin carbons exhibited strong cross peaks with its own aliphatic and hydroxyl protons, e.g. as seen by the T3’4’ and T5,7,9 regions in **Figure 5C**. This has made it impossible to analyze lignin-carbohydrate interactions starting from aromatic carbons. However, clear intermolecular cross peaks were spotted between the well-resolved AcCO and GA6/GlcA6 carbon sites to aromatic protons. Future research should explore 3D DQ-SQ-SQ correlation experiments, incorporating a ^13^C DQ dimension with either two ^1^H dimensions or one additional ^13^C and one ^1^H dimension, as a strategy to further enhance spectral resolution and probe spatial proximities in complex biomolecular systems^89, 90^.

## CONCLUSIONS

This study presents an exploratory application of ^1^H-detected solid-state NMR techniques to investigate the structure and spatial organization of carbohydrates and lignin within intact secondary plant cell walls, using stems from a hardwood (eucalyptus) and a softwood (spruce) as models. We demonstrated that synthesized spectra enable clear identification of key ^1^H resonances corresponding to dominant carbohydrate conformers. The 3D hCCH TOCSY experiment was employed to establish carbon connectivity and assist in completing resonance assignments via ^1^H correlations. By integrating hCCH and hCHH spectra, we effectively resolved carbohydrate–lignin associations, highlighting the contributions of acetylated three-fold xylan conformers—rather than two-fold—as stabilizing molecules at the carbohydrate–lignin interface. In eucalyptus, GlcA sidechains of xylan were found to colocalize with lignin, while cellulose–lignin interactions were revised to involve microfibril surfaces not coated by two-fold xylan. Pectin–lignin interactions were also observed, likely reflecting early-stage lignification. Collectively, these findings provide new molecular insights into the carbohydrate–lignin interface in plant secondary walls. Challenges encountered in this study point to several future directions for improving spectral resolution in lignocellulosic analysis, including partial deuteration, ultrafast MAS, incorporation of double-quantum dimensions into 3D experiments, and improved debarking procedures to minimize tannin interference in softwoods. Beyond the advantages of higher sensitivity, faster acquisition, and enhanced resolution—particularly when combined with ^13^C dimensions—^1^H-detected solid-state NMR also offers strong complementarity with solution-state ^1^H datasets from cell wall extracts, enabling more holistic structural interpretations within the native wall environment.

## Supporting information

Wood Proton SI

## ASSOCIATED CONTENT

Supporting Information. This material is available free of charge via the Internet at http://pubs.acs.org.

Additional planes extracted from 3D hCHH spectrum, ^13^C and ^1^H chemical shifts of carbohydrates and lignin, and experimental conditions.

### Notes

The authors declare no conflict of interest.

## ACKNOWLEDGMENT

This research was supported by the U.S. Department of Energy, Office of Science, Basic Energy Sciences, under award number DE-SC0023702.

## ABBREVIATIONS

CP: cross polarization
DQ: double-quantum
DSS: Sodium trimethylsilylpropanesulfonate
GalA: galacturonic acid
MAS: magic-angle spinning
MISSISSIPI: multiple intense solvent suppression intended for sensitive spectroscopic investigation of protonated proteins
NMR: nuclear magnetic resonance
rf: radio frequency
RFDR: radio frequency-driven recoupling
slpTPPM: swept low power two-pulse phase modulation
SPINAL-64: small phase incremental alteration, with 64 steps
TMS: tetramethylsilane
TOCSY: total correlation spectroscopy
WALTZ-16: wideband alternating-phase low-power technique for zero-residual splitting
Xyl: xylose

