## Supplementary material for "Rapid High-Resolution Analysis of Polysaccharide-Lignin Interactions via Proton-Detected Solid-State NMR with Application to Eucalyptus and Spruce": Wood Proton SI

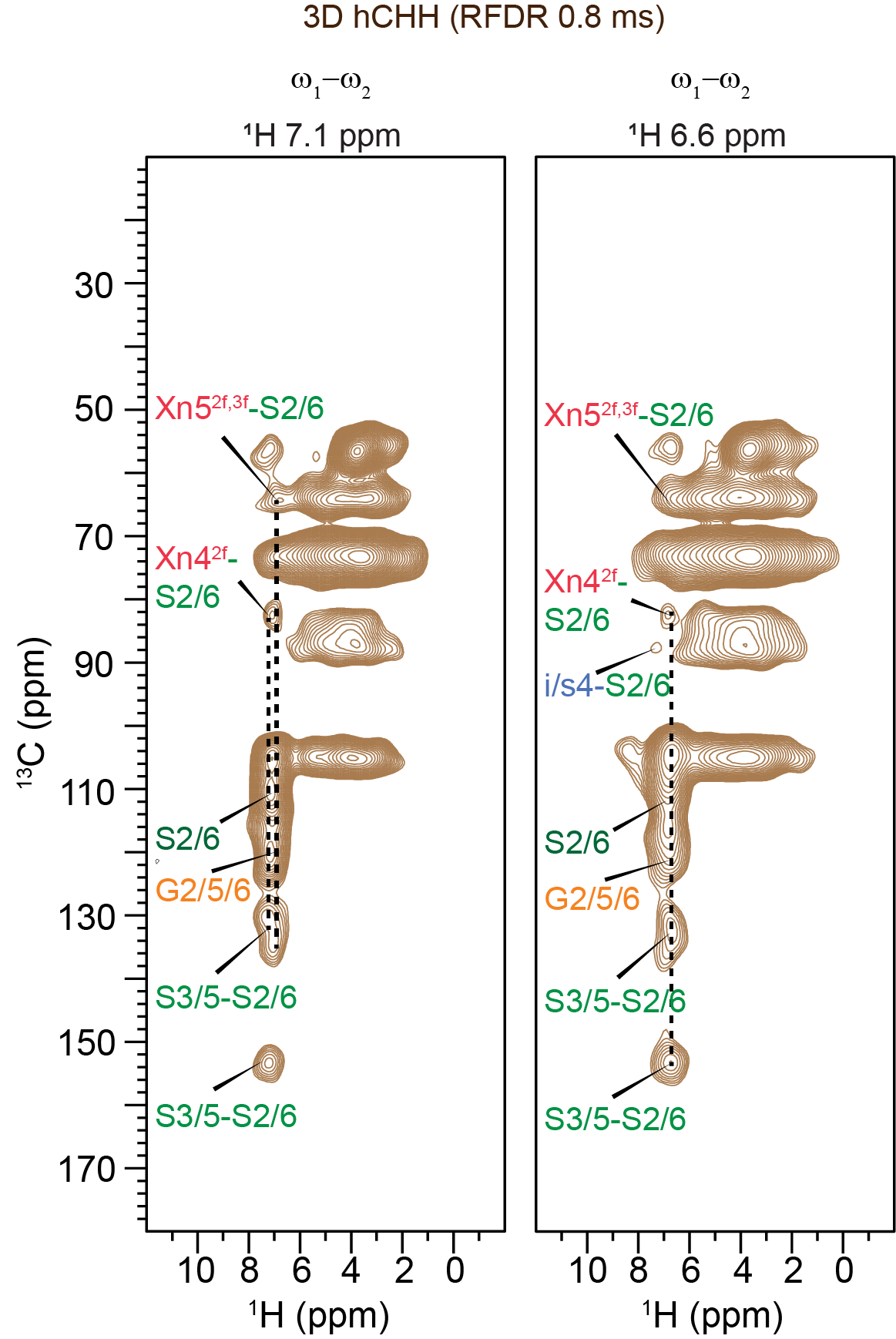

**Figure S1. Through-space lignin-carbohydrates interactions in eucalyptus analyzed by^1^H-^13^C (ω_1_-ω_2_) planes of 3D hCHH spectrum.** 2D strips of ^1^H-^13^C (ω_1_-ω_2_) were extracted at different proton chemical shifts in (ω_3_) from the 3D hCHH RFDR experiment with mixing time of 0.8 ms. The spectra were measured on a 14.1 T spectrometer with a MAS rate of 60 kHz.

**Table S1.  NMR experimental parameters for all samples.** τ_cp1_ and τ_cp2_: the contact times for the first (hC) and the second (Ch and CH) CP, respectively. NS: number of scans; D1 = recycle delay between scans. TD1/2/3 and AQ1/2/3: total data points and chemical shift evolution time for dimensions 1/2/3, respectively.

| Expt. | CP (µs) | | NS | D1  (s) | TD1 | TD2 | TD3 | AQ1  (ms) | AQ2  (ms) | AQ3  (ms) | WALTZ  (ms) | RFDR  (µs) | Expt.  Time | Sample |
| --- | --- | --- | --- | --- | --- | --- | --- | --- | --- | --- | --- | --- | --- | --- |
|  | τ_cp1_ | τ_cp2_ |  |  |  |  |  |  |  |  |  |  |  |  |
| 2D hCH | 1200 | 100 | 16 | 2 | 384  (^13^C) | 2000  (^1^H) | - | 6.4 | 10.0 | - | - | - | 3h39m | Eucalyptus |
|  | 2000 | 50 | 16 | 2 | 384  (^13^C) | 2000  (^1^H) | - | 6.4 | 12.2 | - | - | - | 3h41m |  |
| 2D hChH | 2000 | 500 | 32 | 2 | 1600  (^1^H) | 256  (^13^C) | - | 4.3 | 13.6 | - | - | 133.3  266.7  800.0 | 4h52m |  |
| 3D hCHH | 2000 | 500 | 16 | 2 | 84  (^13^C) | 84  (^1^H) | 1600  (^1^H) | 1.4 | 1.4 | 13.6 | - | 800.0 | 2d18h35m |  |
| 3D hCCH TOCSY | 2000 | 100 | 8 | 2 | 128  (^13^C) | 128  (^13^C) | 1764  (^1^H) | 2.1 | 2.1 | 14.9 | 15 | - | 3d5h51m |  |
| 2D hCH | 1200 | 50 | 8 | 2 | 384  (^13^C) | 2000  (^1^H) | - | 6.4 | 10.0 | - | - | - | 1h49m | Spruce |
|  | 1200 | 1200 | 8 | 2 | 384  (^13^C) | 2000  (^1^H) | - | 6.4 | 12.2 | - | - | - | 1h49m |  |
| 2D CP J-INADEQUATE | 1000 | - | 64 | 1.6 | 340  (^13^C) | 1440  (^13^C) | - | 14.0 | 4.5 | - | - | - | 10h |  |

**Table S2.  ^1^H and ^13^C chemical shifts for the rigid carbohydrates in Eucalyptus.** ^1^H is on DSS scale and ^13^C is on TMS scale. Unidentified sites are indicated as “-”. Not applicable ones are indicated with “/”. Ambiguous assignments (i.e. no previous data to confidently assign to a certain molecule or site) are underlined. The abnormal proton shifts in xlyan are due to potential acetylation and are indicated with “*”.

|  | C1/H1 | C2/H2 | C3/H3 | C4/H4 | C5/H5 | C6/H6 | OMe | AcMe |
| --- | --- | --- | --- | --- | --- | --- | --- | --- |
| i | 105.5/4.58 | 72.2/3.51 | 75.2/3.42 | 88.5/3.28 | 72.2/3.51 | 65.0/3.90 | / | / |
|  | 105.5 | 71.7 | - | 89.0/3.53 | 71.7 | 65.3 | / | / |
|  | 104.6 | - | - | 87.2/3.1 | - | 65.5/3.50 | / | / |
| i/s | 103.2/4.95 | - | - | 86.6 | - | - | / | / |
|  | - | - | - | 86.0/4.38 | - | 65.3 | / | / |
|  | - | - | - | 86.9/4.19 | - | 68.0 | / | / |
| s | 105.4/4.58 | 72.2/3.51 | 75.2/3.42 | 84.8/3.80 | 75.2/3.42 | 61.5/3.72 | / | / |
|  | 104.2/5.12 | 71.1/3.39 | 75.5 | 84.7/2.56 | 75.5 | 62.0/3.34 | / | / |
|  | - | 72.0 | 74.8 | 85.2/3.72 | 74.8 | 64.8/3.70 | / | / |
| Xn2f | 105.0/4.58 | 72.2/3.51 | 75.2/3.42 | 81.8/3.48 | 63.5/3.90 | / | / | 21.2/2.02 |
|  | 105.0 | 73.5 | 74.6 | 82.7/3.72 | 63.6 | / | / | - |
|  | 103.8/5.16* | 72.8/4.02* | 74.4/4.88* | 82.2/4.62* | 62.4/4.02* | / | / | - |
| Xn | - | 73.4/3.90 | 73.4/3.90 | 80.5/3.27 | 64.9/3.62 | / | / | 21.2/2.02 |
|  | - | 73.2 | 73.2 | 80.3/3.92* | 64.2/3.65 | / | / | - |
|  | - | - | - | 80.2/5.1* | 61.1 | / | / | - |
| Xn3f | 102.2/4.58 | 73.3/5.08* | 74.4/4.09* | 78.8/3.63 | 63.9/3.30 | / | / | 21.2/2.02 |
|  | - | 71.34/3.70 |  | 78.3 | 64.3/3.32 | / | / | - |
|  | - | - | 74.2/4.89* | 78.9/3.65 | 65.0 | / | / | - |
|  | - | - | - | 79.2/3.63 | 64.6 | / | / | - |
|  | 102.3 | - | - | 77.5/4.73* | - | / | / | - |
|  | 101.9/5.12 | 71.1 | 73.5 | 77.8 | - | / | / | - |
| Me-GlcA | 98.5/5.30 | 72.7 | 75.7 | 81.5/3.46 | 67.9/4.72 | 176.8 | 59.6/3.58 | / |
|  | 97.8/4.8 | 72.9 | 75.2 | 81.6/4.25 | 66.4/4.4 | 177.6 | - | / |
|  | 98.2 | - | - | 81.8/2.67 | 69.5/3.84 | - | - | / |
|  | 98.1/4.81 | - | - | 83.8 | 70.5 | - | - | / |
|  | 98.1/4.73 | - | - | 80.3 | 70.7 | - | - | / |
| GalA | 100.8/5.14 | 68.0/3.63 | - | 78.5/4.35 | 70.5/3.46 | 171.2 | 53.5/3.75 | 21.2/2.02 |
|  | 101.4/4.81 | 68.8/3.79 | - | 79.0/4.34 | 71.3/3.27 | 170.5 | - | - |
|  | 100.8 | 69.4/4.18 | - | 79.8/4.68 | - | 169.6 | - | - |
|  | 100.8/3.99 | - | - | 75.7 | - | - | - | - |
|  | 99.6/4.7 | 69.9 | - | - | - | - | - | - |
| Rha | - | - | - | - | - | 17.6/1.15 | / | / |
|  | - | - | - | - | - | 19.4/0.91 | / | / |

Note: The OMe, CH_3_ (AcMe and R6) and CO (GalA6 and Me-GlcA6) chemical shifts were identified in 2D hCH and 2D hChH RFDR spectra.

**Table S3.  ^1^H and ^13^C chemical shifts for the rigid lignin in Eucalyptus.** ^1^H is on DSS scale and ^13^C is on TMS scale. Not applicable ones are indicated with “/”. Remote protons are indicated with “*”.

| Lignin site | ^13^C | ^1^H |
| --- | --- | --- |
| OMe | 56.5 | 3.70 |
| S2/6 | 101.5-106.5 | 6.00-7.26 |
| G2  S'2/6 | 109.2 | 6.79 |
|  | 111.2 | 6.98 |
|  | 112.8 | 6.94 |
| G5 | 114.3 | 6.90 |
|  | 115.2 | 6.23/7.31 |
|  | 116.3 | 6.79 |
|  | 117.7 | 7.44 |
| G6 | 119.6 | 7.06 |
|  | 120.3 | 6.03 |
|  | 122.1 | 6.72 |
| S1/G1 | 131.1 | 6.20* |
|  | 132.1 | 6.94* |
|  | 134.1 | 6.72* |
| G3/4 | 147.7 | 7.96* |
| S3/5 | 152.2 | 7.09* |
|  | 152.6 | 6.49* |
|  | 153.1 | 7.65* |
|  | 153.8 | 6.76* |
